# *Wolbachia* promote successful sex with siblings

**DOI:** 10.1101/855635

**Authors:** Zeynab Bagheri, Ali Asghar Talebi, Sassan Asgari, Mohammad Mehrabadi

**Affiliations:** Department of Entomology, Faculty of Agriculture, Tarbiat Modares University, Tehran, Iran; Australian Infectious Disease Research Centre, School of Biological Sciences, The University of Queensland, Brisbane, QLD, Australia

**Keywords:** *Wolbachia*, inbreeding avoidance, *Habrobracon hebetor*, postcopulatory

## Abstract

*Wolbachia* are intracellular α-proteobacteria that have a wide distribution among various arthropods and nematodes. They affect the host reproduction favoring their maternal transmission, which sets up a potential conflict in inbreeding situations when the host avoids sexual reproduction preventing inbreeding depression, while *Wolbachia* pushes it. In this study, we used the wasp *Habrobracon hebetor* to test the hypothesis that *Wolbachia* modulate inbreeding avoidance behaviour and promote sib mating. To test this, we first cured wasps of *Wolbachia* using tetracycline treatment and produced infected and uninfected isolines. Then, we paired the uninfected and infected females with sibling (inbred) and non-sibling (outbred) males in choice and non-choice experiments. Our results showed no obvious precopulatory inbreeding avoidance in this wasp as brother-sister mating rates (in both choice and nonchoice experiments) were not significantly different form non-sibling pairs, regardless of *Wolbachia* infection. However, our results indicated that *H. hebetor* shows a strong postcopulatory inbreeding avoidance behaviour that results in a low fertilization rate of uninfected siblings and therefore high rate of production of male progeny was obtained. We observed higher rates of fertilization success in the *Wolbachia-*infected lines that resulted in significantly higher female progeny production compared to the uninfected sib mates. Since diploid females are the result of successful fertilization due to haplodiploidy sex determination system in this insect, our results indicate that *Wolbachia* promoted fertile sib mating in *H. hebetor.* Interestingly, the rate of adult emergence in the progeny of *Wolbachia*-infected sib mates were almost similar to the non-sib mate crosses and significantly more than those observed in the uninfected sib mate crosses. We support the idea that *Wolbachia* modulate inbreeding avoidance and promote sib mating and also mitigate inbreeding depression. The wasp *Habrobracon hebetor* siblings infected with *Wolbachia* show higher rates of fertilization success and higher adult emergence rates compared to the uninfected sib mates. By promoting successful sex with siblings and increasing the probability of female progeny, *Wolbachia* enhance their transmission to the next generation and also mitigate inbreeding depression. This is an undescribed effect of *Wolbachia* (symbiont) on the host reproduction.

## Introduction

*Wolbachia* are endosymbiotic intracellular bacteria that have a wide distribution in various arthropods, including insects, isopods, and mites, and nematodes (Werren, Baldo & Clark 2008; LePage & Bordenstein 2013). These gram-negative bacteria manipulate the reproductive system of their hosts via male-killing (Jiggins, Hurst & Majerus 1998; Hurst *et al.*, 1999), feminization (Rousset *et al.*, 1992), inducing parthenogenesis (Stouthamer, Luck & Hamilton 1990) and cytoplasmic incompatibility (CI) (Yen & Barr 1971; Turelli & Hoffmann 1995). *Wolbachia* utilize these strategies increasing the frequency of infection in a host population. CI strains of *Wolbachia* cause the diploid eggs obtained from infected males and uninfected females not to hatch (LePage & Bordenstein 2013). In most hosts, CI-*Wolbachia* alter the hosts’ mating behaviour enhancing their occurrence in the population. For example, *Wolbachia* increase the desire for mating in *Drosophila* males increasing the chance of successful mating (De Crespigny, Pitt & Wedell 2006). In addition, infected female mites choose infected males to mate and vice versa (Vala *et al.*, 2004). Thus, in some cases, *Wolbachia* induce reproduction isolation between infected and non-infected populations and create semi-species (Breeuwer & Werren 1990; Bordenstein, O’hara & Werren 2001; Miller, Ehrman & Schneider 2010).

Inbreeding occurs when two closely related organisms mate with each other and produce offspring generally with reduced biological fitness (i.e. inbreeding depression). Inbreeding in wasps including *Habrobracon hebetor* leads to the production of homozygous individuals (diploid males) with lower survival and fertility (Petters & Mettus 1980; Antolin *et al.*, 2003). Because inbred individuals are commonly less fit than outbred individuals due to inbreeding depression, inbreeding avoidance mechanisms at precopulatory (Lihoreau, Zimmer & Rivault 2007; Liu *et al.*, 2014; Pilakouta & Smiseth 2017) and postcopulatory (Tregenza & Wedell 2002; Michalczyk *et al.*, 2011; Duthie, Bocedi & Reid 2016) levels are expected to occur to avoid inbreeding depression. Therefore, female wasps basically avoid mating with their brood-mates preventing inbreeding depression (Ode, Antolin & Strand 1995). This behaviour reduces the mating rate between the siblings, and as a result, male progeny increases. Since *Wolbachia* is maternally inherited, this behaviour could prevent distribution of *Wolbachia*. Therefore, in this study, we questioned whether *Wolbachia* modulate inbreeding avoidance at the pre- or postcopulatory levels in *H. hebetor* in favor of their transmission. If so, what is the fitness consequence of this conflict for the insect host? We show that *Wolbachia* suppress postcopulatory inbreeding avoidance of the female wasps and enhance successful fertilization of the female’s eggs that results in female progeny. This manipulation of the host enhances vertical transmission of *Wolbachia* to the next generations.

## 2. Materials and methods

### 2.1. Insects

The two lab-reared populations of *Habrobracon hebetor* were previously collected from Nazarabad (N) and Mohammadshahr (M), Iran. These populations were naturally infected with *Wolbachia* (W^+^) and reared separately under laboratory conditions at 25 ± 5 °C, 60 ± 5% RH and a photoperiod of 16:8 h, and fed daily with diluted raw honey (90% honey and 10% water). The female wasps were presented with fifth instar Mediterranean flour moth (*Ephestia kuehniella*) larvae for oviposition.

### 2.2. Antibiotic treatment

To test whether *Wolbachia* affect mating behaviour of *H. hebetor* in sib mate and non-sib mate crosses, two isolines from each of the two parasitoid wasp populations were generated (i.e. MW^+^, MW^−^, NW^+^ and NW^−^) using tetracycline treatment (0.2%, w/v with diluted raw honey) as described before (Bagheri *et al.*, 2019). Ten wasps of each isoline was then randomly selected to test for *Wolbachia* infection using qPCR. After successful removal of *Wolbachia* from the wasps (confirmed by qPCR), they were reared for 10 generations without tetracycline treatment under laboratory conditions at 25 ± 5 °C, 60 ± 5% RH and a photoperiod of 16:8 h. The resultant adult wasps from the 13th generation were considered as the uninfected isolines (W^−^) and the adult wasps from these isolines were used for experiments. The *Wolbachia*-infected isolines were reared similarly without antibiotic treatment.

### 2.3. Quantitative PCR (qPCR)

To compare *Wolbachia* density in samples (i.e. MW^+^, MW^−^, NW^+^ and NW^−^), total DNA was extracted from single individuals of *H. hebetor* using a previously described procedure (O’Neill *et al.*, 1992). Concentrations of the DNA samples were measured using an Epoch instrument (BioTek), and 10 ng from each genomic DNA sample was used for qPCR using SYBR Green Mix without ROX (Ampliqon) with a Mic real time PCR (BMS) under the following conditions: 95 °C for 15 min, followed by 45 cycles of 95 °C for 10 sec, 30 sec at the annealing temperature, and 72 °C for 30 sec, followed by the melting curve (72–95 °C). Specific primers targeting the *Wolbachia* cell division protein (*ftsZ*) gene (Kruse *et al.*, 2017) and the insect *RPL27* and *18s rRNA* genes were used as reference genes (Karamipour, Fathipour & Mehrabadi 2016). Reactions from three biological replicates were repeated three times.

### 2.4. Effect of *Wolbachia* on the mating behaviour and fitness of *H. hebetor* under inbred and outbred conditions

The individuals used in the sib mating experiment were derived from one propagation line and thus all individuals were expected to be closely related. In the choice experiments, we allowed females (W^+^ or W^-^) to choose between sib or non-sib males. To do this, 27 virgin females (13 from M and 14 from N propagation lines) were randomly selected and individually placed in a container with a sib and non-sib male and their mate choice behaviours including sib/non-sib mating, number of mating, and mating latency (time to mating) were recorded for 15 min. These experiments were performed for both *Wolbachia* infected and uninfected wasps.

In the non-choice experiments, we randomly selected 21 pairs *Wolbachia* infected isoline (i.e. ♂MW^+^ × ♀MW^+^) and 20 pairs of the uninfected isoline (i.e. ♂MW^−^ × ♀MW^−^), and allowed each pair to mate and lay eggs on the host separately. The rate of mating, mating latency, and mating duration were measured, and the resultant eggs of each pair were separately reared at the abovementioned conditions until the emergence of the adult offspring. Then, 19 pairs of males and females that were obtained from a female i.e. siblings, were mated and allowed to oviposit for 24 hours. The mating rates, numbers of eggs, egg hatching rates, the mean of emerged adults and offspring sex ratio of each pair were recorded for four days. This experiment was performed for *Wolbachia* infected and uninfected wasps.

The individuals used in non-sib mating experiment were derived from two propagation lines (M and N) and thus the individuals were expected to be unrelated. To perform non-sib mating crosses, we collected 100 pupae from the W^+^ and W^-^ isolines from the two propagation lines and allowed them to emerge as female and male adults. Then, 19 virgin females of the M propagation line and 19 virgin males of the N propagation line were mated and allowed to oviposit separately. We calculated the mating rate, the number of eggs, the egg hatching rates, the mean number of emerged adults and the offspring sex ratio for four days.

### 2.5. Data analysis

The mating rates were compared using Mann-Whitney U-test. The mating latency (time to mating), mating duration, mean number of eggs, the rate of hatching eggs, and the percentage of pupae formation, the percentage of progeny emergence and the sex-ratio of progeny emergence per female in different crosses were analyzed by nonparametric methods (Brunner *et al.*, 2002; Shah & Madden 2004) using PROC MIXED procedure of SAS statistical analysis software (SAS).

## 3. Results

To remove *Wolbachia* from *H. hebetor* individuals, they were treated with tetracycline and our qPCR results confirmed that *Wolbachia* had been successfully removed from tetracycline-treated wasps (Fig. S1). To determine the effect of *Wolbachia* on the mating behaviour of the wasp, we performed mate choice tests. The results of female mate choice experiments showed no significant differences in the mating rate and mating latency (time to mating) between sib and non-sib males or mating latency in general, regardless of *Wolbachia* infection status (Fig. 1A-C, Table 1).

**Table 1.**
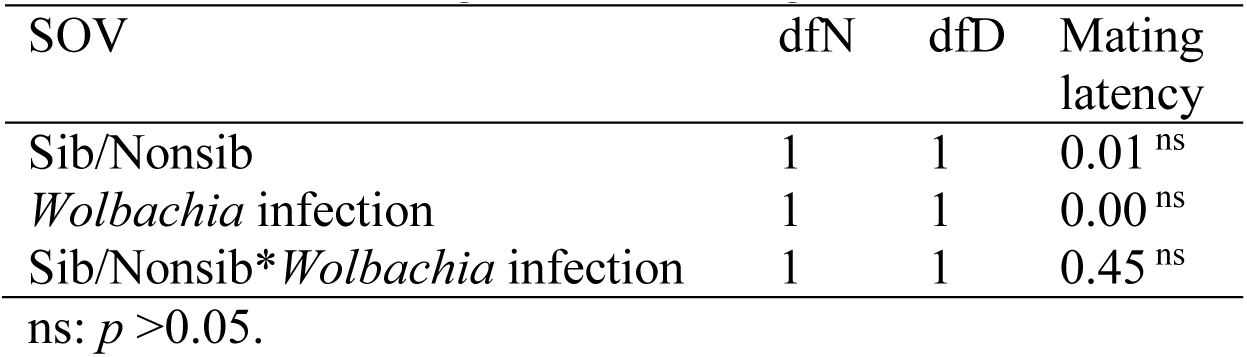
Analysis of variance (Chi-square) for the effect of *Wolbachia* on *H. hebetor* mating behaviour with siblings and nonsibling.

| SOV | dfN | dfD | Mating latency |
| --- | --- | --- | --- |
| Sib/Nonsib | 1 | 1 | 0.01<sup>ns</sup> |
| *Wolbachia* infection | 1 | 1 | 0.00<sup>ns</sup> |
| Sib/Nonsib* *Wolbachia* infection | 1 | 1 | 0.45<sup>ns</sup> |

ns: $p > 0.05$ .

**Fig. 1.**
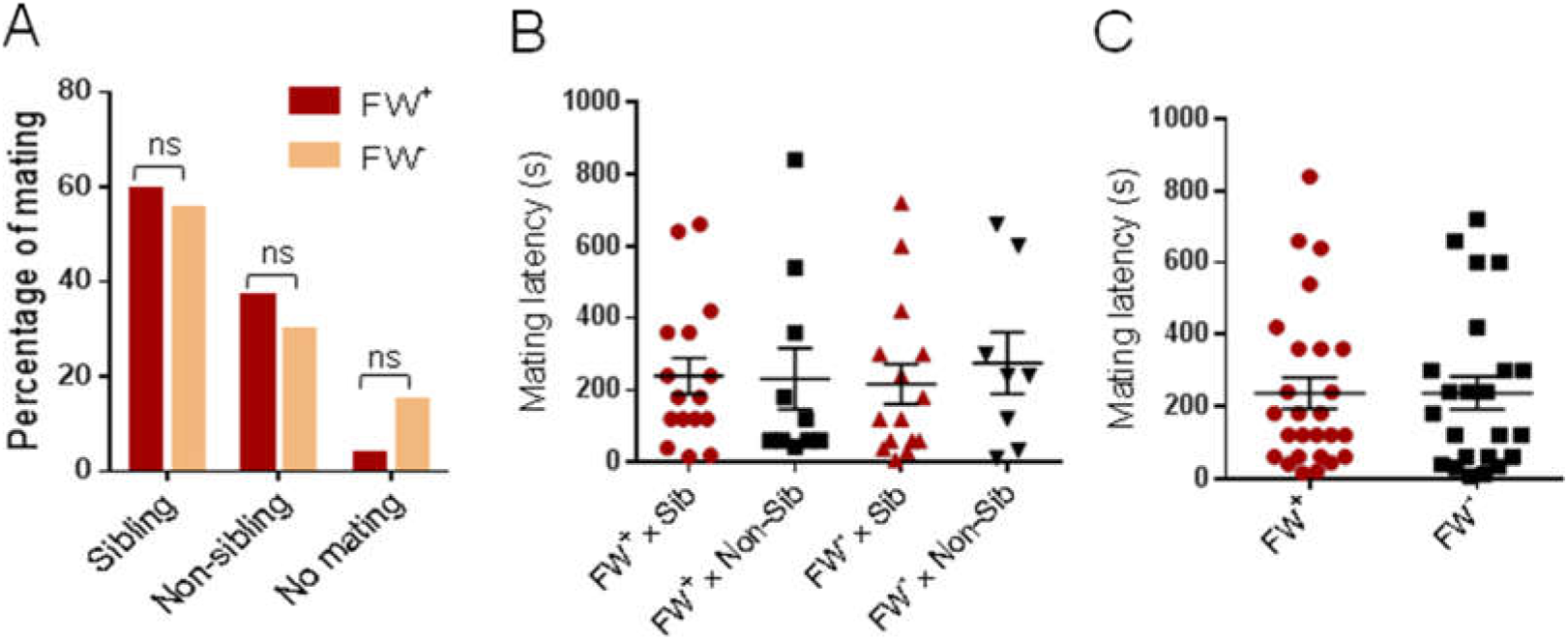
No evidence for precopulatory inbreeding avoidance in *Wolbachia*-infected and uninfected *H. hebetor*. A) Percentage of female mating with sib and non-sib males (choice mating) in the *Wolbachia*-infected and uninfected wasps. B) Time to female mating (mating latency) with sib and non-sib males. C) Mating latency of *Wolbachia*-infected and uninfected females with males. *Wolbachia*-infected (FW^+^) and uninfected (FW^-^) females. Mann-Whitney *U* test.

To investigate the effect of *Wolbachia* on the reproduction and fitness of *H. hebetor* in sib mating (non-choice), we created four different crosses including *Wolbachia-*infected sibling pairs, uninfected sibling pairs, *Wolbachia*-infected non-sibling pairs and uninfected non-sibling pairs, and analyzed different parameters associated with reproduction of the wasps. The results showed no differences in the mating latency, attempts to mate, and mating duration between the different crosses (Fig. 2A-C, Table 2). However, the results showed that the highest numbers of female progeny per female were generated in the progeny of *Wolbachia*-infected non-sibling and uninfected non-sibling pairs (63.9 ± 2.28 and 52.59 ± 4.38, respectively). Interestingly, the *Wolbachia-*infected sibling pairs produced higher female progeny per female compared to uninfected sibling pairs (27.8 ± 6.89 and 9.04 ± 3.32 respectively). In fact, the uninfected sibling pairs produced the lowest number of female progeny (Fig. 2D. Table 2).

**Table 2.**
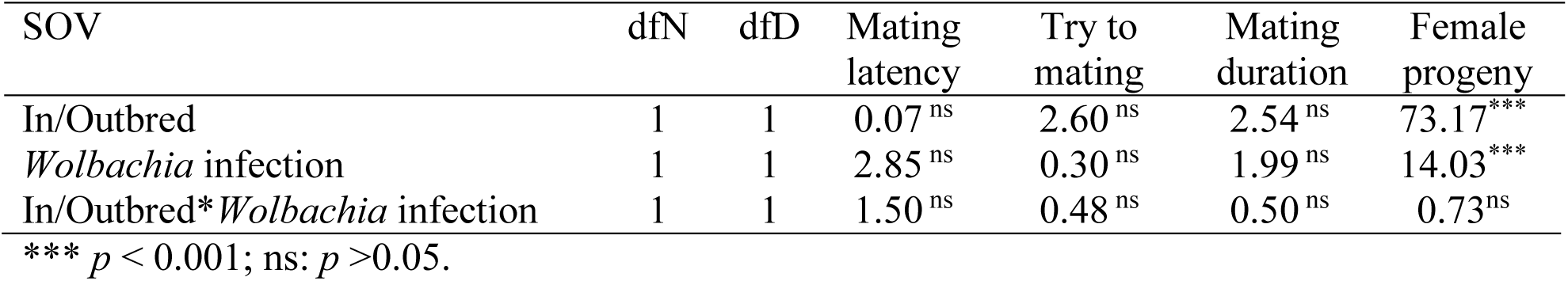
Analysis of variance (Chi-square) for the effect of *Wolbachia* on inbreeding avoidance behaviour of *H. hebetor.*

| SOV | dfN | dfD | Mating latency | Try to mating | Mating duration | Female progeny |
| --- | --- | --- | --- | --- | --- | --- |
| In/Outbred | 1 | 1 | 0.07<sup>ns</sup> | 2.60<sup>ns</sup> | 2.54<sup>ns</sup> | 73.17<sup>***</sup> |
| *Wolbachia* infection | 1 | 1 | 2.85<sup>ns</sup> | 0.30<sup>ns</sup> | 1.99<sup>ns</sup> | 14.03<sup>***</sup> |
| In/Outbred* *Wolbachia* infection | 1 | 1 | 1.50<sup>ns</sup> | 0.48<sup>ns</sup> | 0.50<sup>ns</sup> | 0.73<sup>ns</sup> |

\*\*\* $p < 0.001$ ; ns: $p > 0.05$ .

**Fig. 2.**
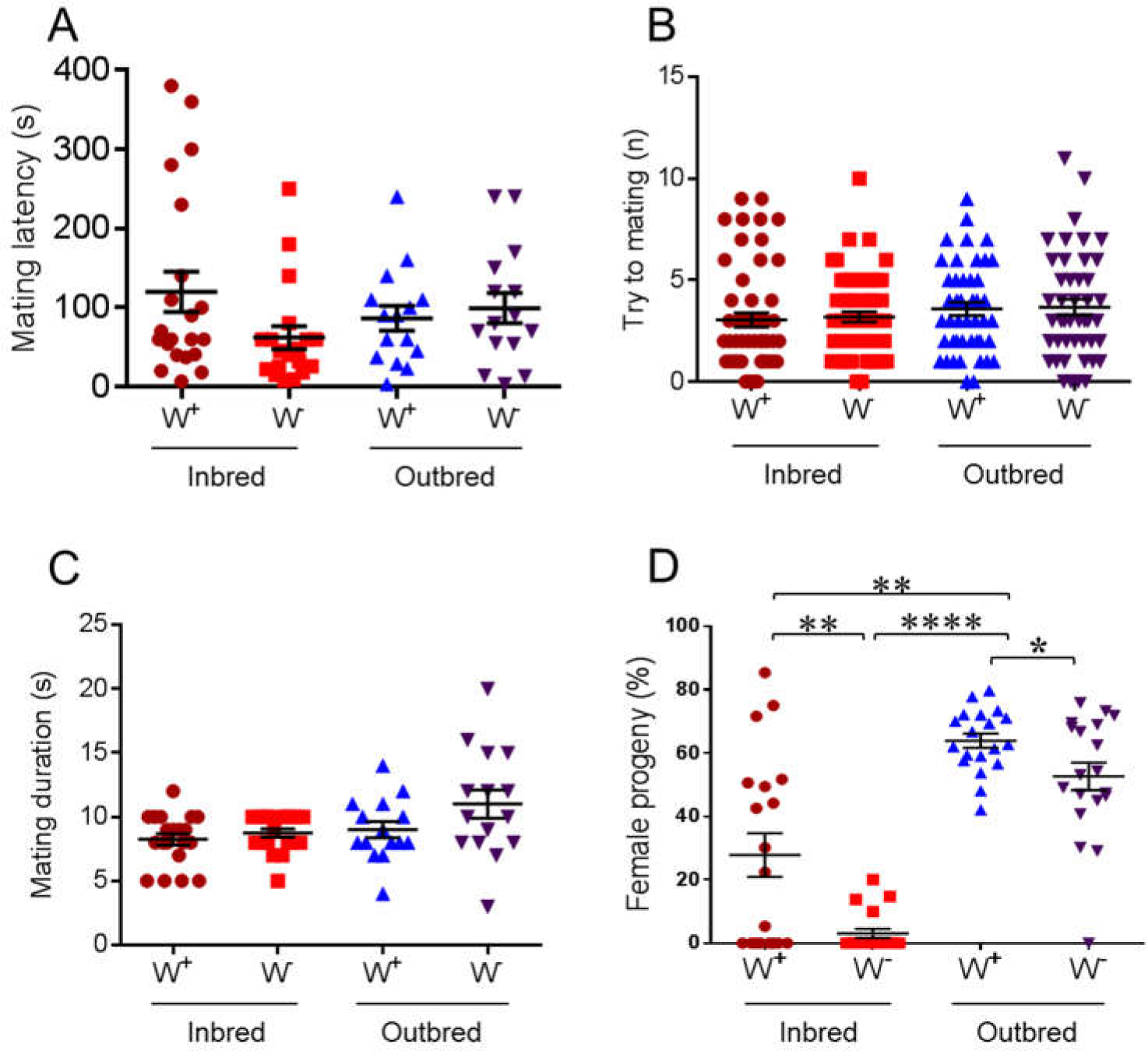
Evidence for postcopulatory inbreeding avoidance in *H. hebetor* and influence of *Wolbachia* on it. A) Time to female mating with sib and non-sib males (non-choice mating) in the *Wolbachia*-infected and uninfected wasps. B) The number of male try to mate with females in the *Wolbachia*-infected and uninfected wasps. C) Siblings and non-siblings mating duration in the *Wolbachia*-infected and uninfected wasps. D) Percentage of female progeny in the inbred and outbred lines of the wasp in the presence and absence of *Wolbachia*. Each data point represents the percentage of female offspring produced per female during four days. *Wolbachia*-infected (W^+^) and uninfected (W^-^). Mann-Whitney *U* test, *, *p* < 0.05; **, *p* <0.01; ***, *p* <0.001.

We also compared the number of resultant eggs of the crosses during four days of oviposition. The mean numbers of eggs within four days per female of *Wolbachia*-infected non-sibling pairs were lower than others (Fig. 3A, Table 3). In addition, the rates of egg hatching in sibling pairs were higher than non-sibling pairs; moreover, the rates of egg hatching in the uninfected crosses were higher than the infected crosses (Fig. 3B, Table 3). The lowest rate of pupa formation was observed in the uninfected non-sibling (91.09±1.21), while there was no difference among others (Fig. 3C, Table 3). The progeny emergence rate of uninfected sibling pairs was significantly lower than that of other crosses (55.16±3.40) including *Wolbachia*-infected sibling pairs. The adult progeny emergence rate of the *Wolbachia*-infected sibling pairs was similar to those of non-sibling crosses (Fig. 3D).

**Table 3.** Analysis of variance (Chi-square) for the effect of *Wolbachia* on biological parameters of *H. hebetor* in different crosses.

| SOV | dfN | dfD | Egg | Hatching | Pupa | Adult |
| --- | --- | --- | --- | --- | --- | --- |
| In/Outbred | 1 | 1 | 9.00<sup>**</sup> | 62.54<sup>***</sup> | 22.06<sup>***</sup> | 43.60<sup>***</sup> |
| *Wolbachia* infection | 1 | 1 | 0.88<sup>ns</sup> | 13.3<sup>***</sup> | 7.33<sup>***</sup> | 24.05<sup>***</sup> |
| In/Outbred* *Wolbachia* infection | 1 | 1 | 0.51<sup>ns</sup> | 0.09<sup>ns</sup> | 0.07<sup>ns</sup> | 9.18<sup>**</sup> |

\*\*\* $p < 0.001$ ; \*\* $p < 0.01$ ; ns: $p > 0.05$ .

**Fig.3.**
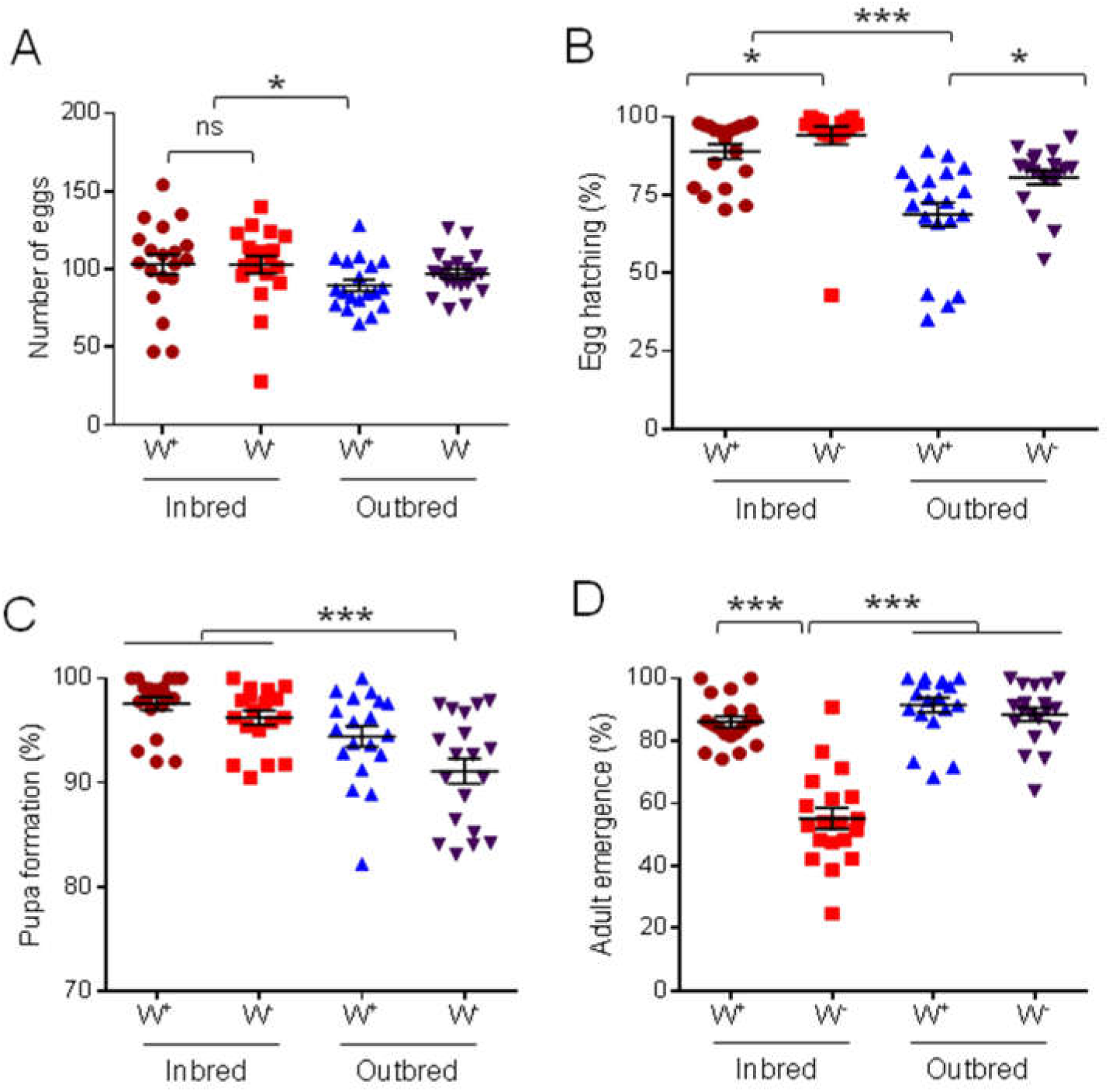
Comparison of biological parameters of *H. hebetor* in different crosses. A) Comparison of the mean number of eggs per female; B) The egg hatching rate; C) The pupa formation rate; D) The percentage of progeny emergence in different crosses. *Wolbachia*-infected (W^+^) and uninfected (W^-^). Each data point represents a sum of data obtained per female during four days. Mann-Whitney *U* test, *, *p* < 0.05; **, *p* <0.01; ***, *p* <0.001.

## 4. Discussion

Inbreeding avoidance has been seen in many species, to avoid mating with siblings (Lihoreau, Zimmer & Rivault 2007; Michalczyk *et al.*, 2011; Liu *et al.*, 2014; Duthie, Bocedi & Reid 2016; Pilakouta & Smiseth 2017). The progeny resulting from inbreeding have less survival, thus females are less willing to mate with siblings than non-siblings (Harper *et al.*, 2016). We previously showed that *Wolbachia* induce a strong CI in *H. hebetor* and affect its mating behaviour (Bagheri *et al.*, 2019). In this study, we investigated the effect of *Wolbachia* on pre- and postcopulatory inbreeding avoidance behaviour in the parasitoid wasp, *H. hebetor*. Further, we determined the effect of *Wolbachia* on the offspring sex-ratio from *H. hebetor* in the sib/non-sib mating conditions.

Our results showed that despite no differences in mating rate and mating latency in sib and non-sib mating crosses (choice and nonchoice female mating), higher percentages of females were produced in the non-sib mating crosses compared to the sib mating crosses suggesting the existence of a postcopulatory inbreeding avoidance mechanism in this parasitoid wasp. However, in *Wolbachia*-infected sib mating crosses, we observed higher percentages of females compared to the uninfected sib mating crosses, while the rates of mating and mating latency were not different. Considering females are the result of sexual reproduction and successful fertilization in the wasp, it can be concluded that inbreeding avoidance is suppressed in the presence of *Wolbachia* and as a result, infected female wasps more readily allow sibling male’s sperms to fertilize their eggs. In other words, it seems that *Wolbachia* suppress postcopulatory inbreeding avoidance and modulate the female to fertilize its eggs with sperms from siblings thereby increasing the number of female progenies that transmit it to the next generation. There is no previous report on the effect of *Wolbachia* on the inbreeding avoidance, however, it has been shown that *Wolbachia* can affect male fertility and sperm characteristics. In *D. simulans*, for example, it has been reported that *Wolbachia*-infected males mate at a higher rate than uninfected males (Awrahman, Champion De Crespigny & Wedell 2014). Moreover, *Wolbachia* infection has been shown to reduce sperm competitive ability in *D. simulans* (Champion De Crespigny & Wedell 2006). Lower rate of sperm transfer in *Wolbachia*-infected males compared to uninfected males has also been reported in *D. simulans* (Awrahman, Champion De Crespigny & Wedell 2014) and the Mediterranean flour moth, *Ephestia kuehniella* (Lewis *et al.*, 2010). Our results beside these reports show that *Wolbachia* infection is able to affect the host reproductive strategies that have important consequences on *Wolbachia* transmission and population dynamics.

In addition, we investigated the effect of *Wolbachia* on the egg numbers, egg hatching rate, pupa formation and the adult emergence rate of *H. hebetor* in sib/non-sib mate crosses. The results showed that the egg numbers, egg hatching and pupa formation rates were significantly higher in sib mate crosses in comparison to non-sib mates regardless of *Wolbachia* infection status. However, we observed significantly lower rates of adult emergence in uninfected sib mates. In fact, the rate of adult emergence in *Wolbachia*-infected sib mates were similar to non-sib mate crosses. These results showed that *Wolbachia* infection mitigated the effect of inbreeding on the adult emergence rate in the progeny of sib mates. It has been reported that inbreeding reduces fitness of offspring especially their survival rate (Meunier & Kölliker 2013; Pilakouta & Smiseth 2016). For example “*Plutella xylostella*” and “*Phaedon cochleariae*” suffered from inbreeding depression; egg hatching rate, survival, fecundity and demographic parameters of the inbred line significantly declined compared to those of the outbred line over time (Peng *et al.*, 2015; Müller, Lamprecht & Schrieber 2018). However, our data suggests that *Wolbachia* mitigated fitness depression raised from sib mating.

This study is the first to show that a symbiont (i.e. *Wolbachia*) interferes with inbreeding avoidance of its host post-copulatory and promote successful mating with siblings. This reproductive modification increases the number of female offspring that could vertically transmit *Wolbachia* to the next generations. We also support the idea that *Wolbachia* alleviate inbreeding depression effects in inbred lines highlighting a complex host modification by this bacterium to increasing its probability of transmission to the next generation.

## Acknowledgments

We thank Prof. Ary Hoffmann (University of Melbourne) and Prof. David Hosken (University of Exeter) for their insightful comments and suggestions. Authors declare no competing interests. All data is available in the main text or the supplementary materials.

## Authors’ contribution

Z.B. and M.M. designed the experiments. Z.B. performed the experiments. Z.B., M.M., A.A.T., S.A. analyzed the data and wrote the paper.

## Data accessibility statement

should the manuscript be accepted, the data supporting the results will be archived in an appropriate public repository (Dryad) and the data DOI will be included at the end of the article.

---

**Fig. S1.**
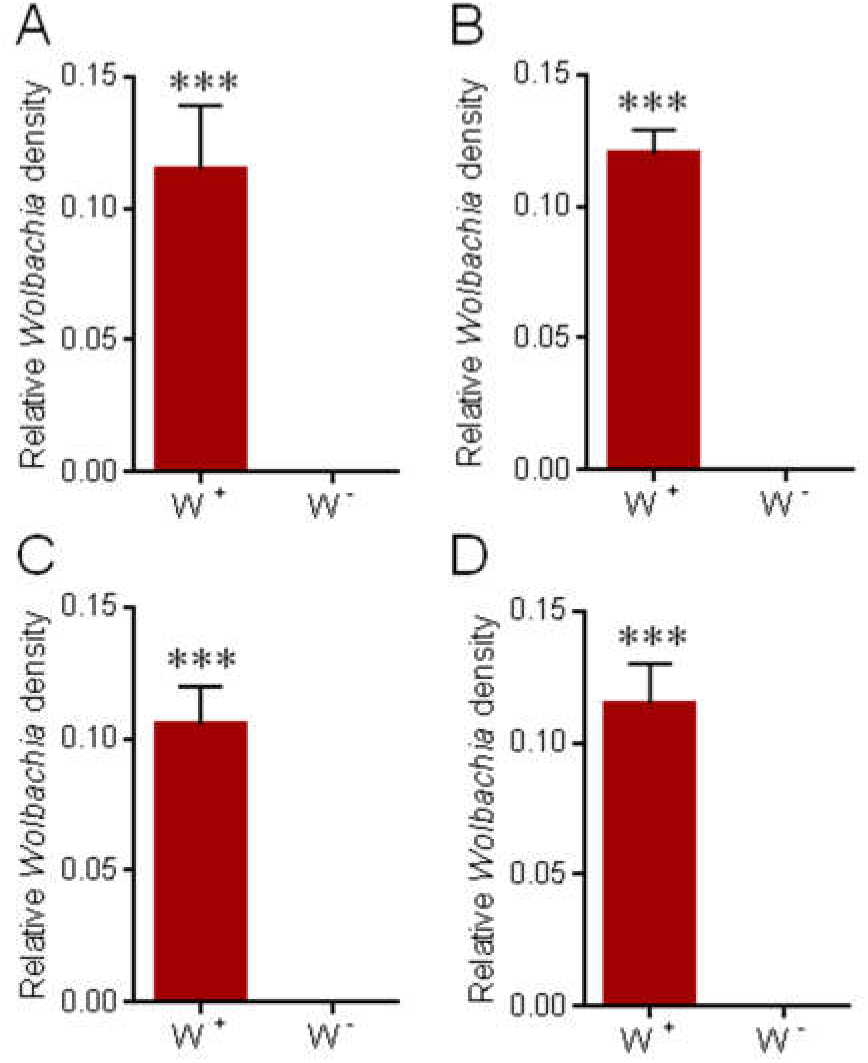
Comparison of *Wolbachia* density using qPCR analysis. QPCR analysis of DNA extracted from *Wolbachia*-infected (W^+^) and uninfected (W^-^) isolines of parasitoid wasps from Nazarabad (A, B) and Mohammadshahr (C, D) Karaj population using tetracycline treatment. Primers to the *Wolbachia ftsZ* gene were used. ***, *p* <0.001.

